# Use of whole genome sequencing to investigate an outbreak of gonorrhoea among females in urban New South Wales, Australia, 2012 to 2014

**DOI:** 10.1101/175869

**Authors:** Cameron Buckley, Brian M. Forde, Ella Trembizki, Monica M. Lahra, Scott A. Beatson, David M. Whiley

## Abstract

Increasing rates of gonorrhoea have been observed among urban heterosexuals within the Australian state of New South Wales (NSW). Here, we applied whole genome sequencing (WGS) to better understand transmission dynamics. Ninety-four isolates of a particular *N. gonorrhoeae* genotype (G122) associated with female patients (years 2012 to 2014) underwent phylogenetic analysis using core single nucleotide polymorphisms (SNPs). Context for genetic variation was provided by including an unbiased selection of 1,870 *N. gonorrhoeae* genomes from a recent United Kingdom (UK) study. NSW genomes formed a single clade, with the majority of isolates belonging to one of five clusters, and comprised patients of varying age groups. Intra-patient variability was less than 7 core SNPs. Several patients had indistinguishable core SNPs, suggesting a common infection source. These data have provided an enhanced understanding of transmission of *N. gonorrhoeae* among urban heterosexuals in NSW, Australia, and highlight the value of using WGS in *N. gonorrhoeae* outbreak investigations.

## Background

Notification rates for *Neisseria gonorrhoeae* infection in Australia have almost doubled in the last 5 years (1). Gonorrhoea is primarily concentrated in two key populations in Australia: men who have sex with men (MSM) living in urban areas and Indigenous heterosexuals in regional and remote settings (2). Until recently, there have been relatively low rates of gonorrhoea among heterosexuals living in urban areas of Australia (3, 4). A retrospective study involving two sexual health clinics in urban Sydney, New South Wales (NSW), showed increasing rates among heterosexuals from 2008 to 2012 (3) that were associated with unprotected oral sex and commercial sex work (CSW). Our subsequent nationwide study of *N. gonorrhoeae* genotypes conducted using isolates from throughout Australia in 2012 revealed the most common gonococcal strain was associated with heterosexuals, and was particularly prevalent among females in NSW (5). A further investigation (6) showed this strain remained prevalent within NSW in 2014 and, although not remarked upon in the article, was prevalent among females. Combined, the above studies are highly suggestive of an emerging epidemic of gonorrhoea among urban heterosexuals in Australia.

A limitation of the genotyping protocol used in our previous studies was that whilst it targets 25 informative single nucleotide polymorphisms (SNPs) across the gonococcal genome, it lacks the resolution offered by whole genome sequencing (WGS). The limitations of traditional genotyping methods have been highlighted by several *N. gonorrhoeae* sequencing studies (7, 8). WGS now has the ability to identify *N. gonorrhoeae* transmission links in the absence of traditional contact-tracing data (8), as well as to reveal geographical and temporal spread and bridging across sexual networks of particular strains (9).

In this study we aimed to utilise WGS to better understand transmission dynamics within a potentially new outbreak of *N. gonorrhoeae* infection among heterosexuals living within urban areas of NSW, Australia. This was prompted by the above studies and more recent unpublished data showing increasing notifications among older individuals in urban areas of Australia, including females. Understanding these trends is important, particularly in light of the potential for increasing disease, including pelvic inflammatory disease among females and also because older age is not typically viewed as a key risk factor for acquiring gonorrhoea amongst heterosexuals. Here, we sequenced the genomes of all available isolates from females that belonged to the most common gonococcal strain identified in the previous studies (5, 6).

## Methods

### *N. gonorrhoeae* isolates

A total of 94 *N. gonorrhoeae* isolates from 87 female patients from two previous investigations were included in this study. The samples comprised 31 vaginal swabs, 27 cervical swabs, 33 throat swabs, 1 rectal swab, 1 eye swab and 1 genital swab. Age groups for the patients were; 18-24 years (n = 29), 25-34 (n = 31), 35-44 (n = 14), 45-54 (n = 11), 55 and over (n = 9). The isolates were from NSW, Australia and were isolated in January to June of 2012 (n = 59) and January to June of 2014 (n = 35). The isolate metadata is summarised in Supplementary Table 1. All 94 isolates were indistinguishable on the basis of iPLEX massarray genotyping. It should be noted that different names were used to describe these strains in the previous studies; the year 2012 isolates were originally described as belonging to strain-type “S01” and genotype “G122” (Supplementary Table 3 in (5)). The year 2014 isolates were called “NSW-001”. For consistency in this study, all 94 isolates will be referred to as being of genotype G122.

### *N. gonorrhoeae* isolates from the United Kingdom

To provide context for the degree of variation among the NSW genomes, analyses of genome data from 1,870 *N. gonorrhoeae* isolates from a recent investigation (8) based in Brighton, United Kingdom (UK) were included. The UK genome data was chosen for comparison because it was the largest collection of *N. gonorrhoeae* genome data with unbiased isolate selection and a collection period (2011-2015) that coincided with the NSW isolates (collected in 2012 and 2014).

### DNA extraction, sequencing and assembly

A single colony from each isolate was cultured on LB agar and incubated at 37^o^C, enriched with 5% CO_2_ for approximately 18-24 hours. Genomic DNA was extracted from half of a 10µl loop of culture growth using the Ultraclean Microbial DNA Isolate Kit (GeneWorks, Australia), using the ‘Alternative Lysis Method’ as per manufacturer’s instructions. Libraries were prepared using the Illumina Nextera XT protocol, with 125bp paired-end reads generated using the Illumina HiSeq instrument (AGRF, Melbourne, Australia), with HiSeq 2500 V4 chemistry. Raw sequence reads from the 94 samples were assessed in FastQC v0.11.4 (10) and hard trimmed to 100bp using Nesoni v0.132 (11). The 94 NSW genomes were assembled in parallel with the 1,870 *N. gonorrhoeae* genomes from the UK (8). The UK genomes were acquired from the NCBI Short Read Archive under BioProject PRJNA315363 (https://www.ncbi.nlm.nih.gov/bioproject/315363). The quality trimmed reads from both NSW and the UK datasets underwent *de novo* assembly using SPAdes v3.6.2 and MegaHit v1.0.3 (13), respectively. Sequence read and assembly metrics are summarised for all NSW genomes in Supplementary Table 2. Sequence read data for all NSW isolates were submitted to the NCBI Short Read Archive under BioProject PRJNA392203 (https://www.ncbi.nlm.nih.gov/bioproject/392203). The taxonomic sequence classification tool Kraken v0.10.5 (14) was initially used to confirm each genome as *N. gonorrhoeae* species. Exclusion criteria applied to the UK genomes included: (i) a genome size greater than 3Mbp, (ii) a genome that had >350 contigs and, (iii) a genome percentage identity according to Kraken of <75% for *N. gonorrhoeae*.

### Genotyping

*In silico* analyses were used to determine multilocus sequence type (MLST) for all NSW and UK genomes. Briefly, the MLST profile was determined using mlst v2.1 (15), providing a seven-locus typing scheme commonly used for Neisseria species (*abcZ, adk, aroE, fumC, gdh, pdhC* and *pgm*). For the 94 NSW genomes, *N. gonorrhoeae* multi-antigen sequence type (NG-MAST) was determined using NGMASTER (16). Sequences with different NG-MAST profiles were compared using BioEdit v7.0.9.0 (17). Novel NG-MAST profiles were uploaded to the NG-MAST database (http://www.ng-mast.net/). The NG-MAST predictions for the UK genomes were kindly provided by Dr David Eyre, University of Oxford, UK (personal communication).

### Phylogenetic analyses

All phylogenetic trees were constructed using a core SNP alignment following the removal of recombinant regions. Genomes were aligned using Parsnp v1.2 (18) by randomly selecting a NSW genome (AU2012-573) to act as a reference, followed by the prediction and removal of recombinant regions using Gubbins v2.1.0 (19). Phylogenetic analyses based on core SNP alignments for both the NSW and UK genomes were achieved using RAxML v8.2.9 (20) using a general-time reversible nucleotide substitution model with a GAMMA correction for site variation. Phylogenetic trees were visualised using FigTree v1.4.2, Evolview v2 or iTOL v3 (21-23).

### Mutations associated with antimicrobial resistance

All 94 NSW genomes and closely related UK genomes, underwent annotation via Prokka v1.11 (24). A combination of the method outlined below and BIGSdb (25) were used to assess loci associated with antimicrobial resistance (AMR) including; *gyrA*, *folP*, *mtrR*, *parC*, *parE*, *penA*, *ponA*, *porB*, *rpoB*, *rpsE*, *rspJ* and 23S rRNA. Using BLAST+ and reference sequences derived from characterised WHO-F and WHO-U strains (26), loci were extracted from an annotated multi-fasta file using seqtk (27). Sequences were aligned and inspected using Clustal Omega (28).

## Results

### Phylogenetic analysis of NSW genotype G122 isolates in the context of a broad collection of UK isolates

The core SNP phylogeny (Figure 1A) included all NSW isolates and 1,796 of the 1,870 UK genomes; 74 UK genomes were removed from further analysis on the basis of the above exclusion criteria. The unrooted maximum likelihood phylogeny in Figure 1A shows a diverse range of strains from the UK. Figure 1B shows all 94 NSW genomes tightly cluster in a single clade with six UK genomes (SRR3350090, SRR3343655, SRR3350214, SRR3350225 and SRR3343534, SRR3350146). All genomes in this clade (hereafter called the G122 clade) were MLST 7359, while all 6 UK genomes and the majority of NSW genomes (n = 86) were NG-MAST 4186. The remaining NSW isolates comprised four previously described NG-MAST types (NG-MAST 6759, n = 2; NG-MAST 6767, n = 2; NG-MAST 15344, n = 2 and NG-MAST 15348, n = 1) as well as one novel NG-MAST profile (NG-MAST 15609, n = 1). All NG-MAST profiles shared the same *tbpB* 241 allele but differed by their *porB* sequences which shared >99% nucleotide identity, indicating they all belonged to the same genogroup (29); which we assigned as “genogroup 4186”. Based on the most recent common ancestor, isolates in the G122 clade differ by <200 core SNPs from isolates in the nearest sister clade, comprising 9 UK genomes that are all MLST 7826 and NG-MAST 2487 (Figure 1B). Both clades form a distinct lineage approximately 1000 core SNPs distant to the nearest split in the underlying *N. gonorrhoeae* phylogeny (Figure 1A).

**Figure 1.**
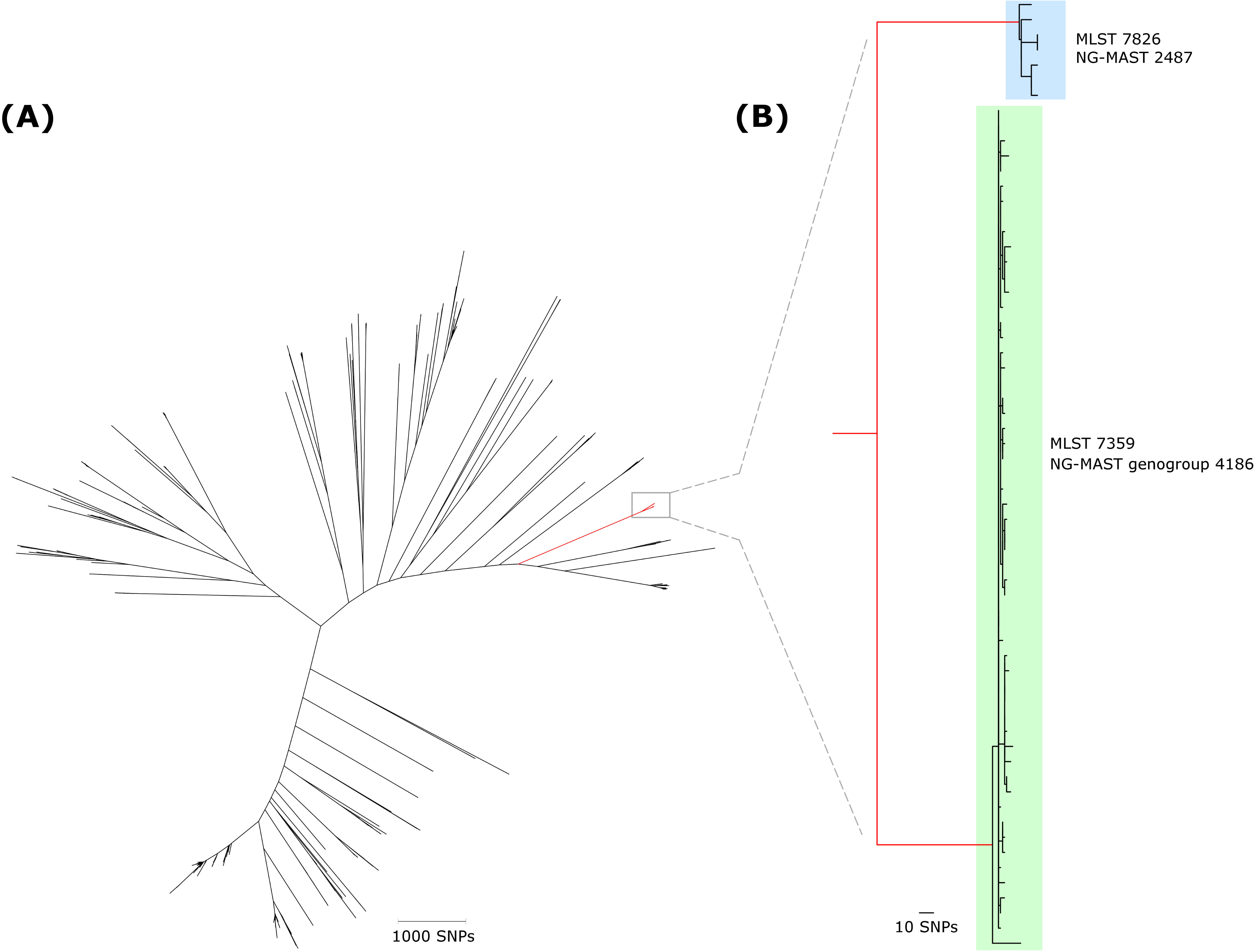
**(A)** An unrooted maximum likelihood phylogeny constructed with 14,802 core single nucleotide polymorphisms (SNPs) from both New South Wales (NSW; n = 94) and the United Kingdom (UK; n = 1,796) genomes based on non-recombinant core regions. The scale bar represents SNPs. **(B)** An enhanced view of the branch containing two clades highlighted in green and blue. The green clade includes all NSW genomes and 6 UK genomes which comprised MLST 7359 and NG-MAST genogroup 4186. The next closest sister clade (in blue) comprised 9 unrelated MLST (7826) and NG-MAST (2487) UK genomes. The scale bar represents SNPs.

### Phylogenetic analysis of the G122 clade

To investigate the genetic relationships between G122 isolates at the single nucleotide level we determined a core SNP phylogeny of all 94 NSW genomes and the six related UK genomes (Figure 2). The majority of isolates belong to one of five clusters (C1 to C5) comprising six or more isolates that each share a defining SNP and a common ancestor indicating a transmission network (Figure 2). The maximum core SNP difference between genomes in each cluster, C1 to C5 respectively, were 6, 4, 19, 21 and 14 SNPs. Of these clusters, only three (C3, C4 and C5) comprised isolates from both years 2012 and 2014 whereas C1 and C2 comprised isolates from year 2012 only. Age group for clusters C3 to C5 ranged from 18-24 to 55 and over (median = 25.5, 32 and 25 years respectively), while C1 ranged from 18-24 to 45-54 (median = 35.5) and C2 ranged from 25-34 to 45-54 (median = 45.5). The percentage of cervical/vaginal and throat samples respectively for each cluster were 33% and 67% for both C1 and C2, 57% and 36% for C3, 65% and 35% for C4 and 58% and 42% for C5 (summarised in Supplementary Table 3).

**Figure 2.**
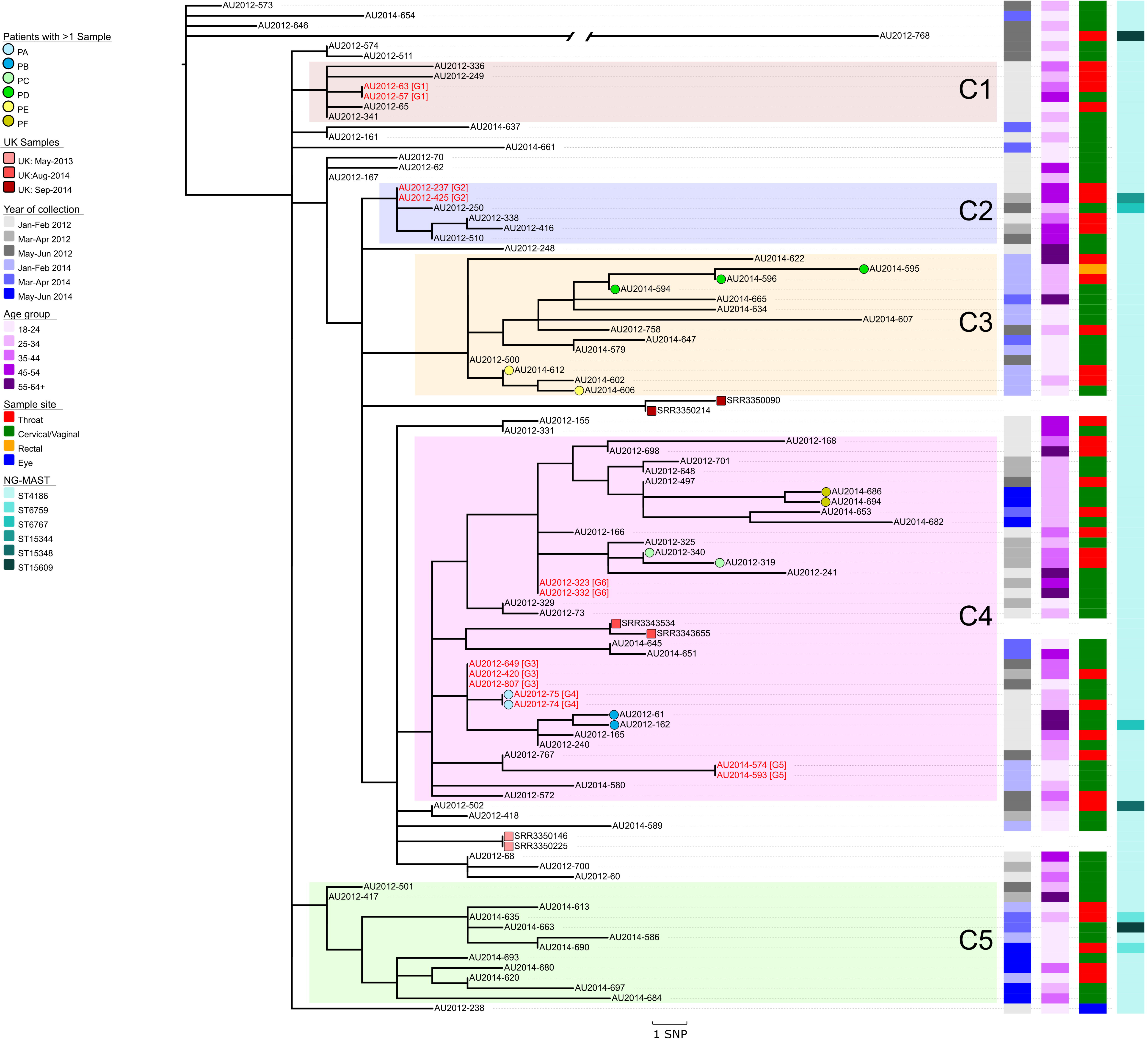
A rooted maximum likelihood phylogeny constructed with 473 core single nucleotide polymorphisms (SNPs). It comprises the G122 clade containing all 94 New South Wales (NSW) genomes and 6 United Kingdom (UK) genomes. A UK genome (SRR3360636) from the next closest sister clade was selected as the outlying group to root the phylogeny. Five larger clusters of isolates (C1-C5) have been labelled based on the phylogeny. Coloured circles represent isolates from the same patient and coloured squares represent isolates from the UK (see key). Rectangles and their respective colour schemes correspond with date of collection, age group, sample site and NG-MAST profile (see key). The red isolate names with G1-G6 appended to the end, represent isolates that share identical core SNPs. Genome AU2012-768 has 33 core SNPs however its branch is truncated for easier visualisation purposes. The scale bar represents SNPs.

Genome AU2012-768 contained 33 distinct core SNPs, recombination analysis identified three highly SNP-dense regions that were filtered prior to tree-building. Manual inspection of these regions revealed multiple SNPs were shared between AU2012-768 and the outlying UK genome (SRR3360636) used to root the phylogeny.

### Diversity of G122 isolates within patients

Six individuals with more than one isolate were included in our study (Patients ‘PA’ to ‘PF’). The respective genomes from these six patients differed by less than seven core SNPs. Patient ‘PA’ had a throat and cervical swab collected on the same day, and was the only individual with more than one isolate that had indistinguishable core SNPs. Patients ‘PB’, ‘PC’ and ‘PD’ also had their respective isolates collected on the same day from different sample sites, but showed a difference of 2-7 core SNPs. The remaining patients, PE and PF, each had different isolates collected between one and eleven days respectively and both differed by 2 core SNPs.

### Genetic links between patients

There were 6 instances where isolates had indistinguishable core SNPs and have been labelled as groups G1 to G6; these comprised either 3 isolates (G3) or 2 isolates each (G1, G2, G4, G5 and G6). Group G4 involved a single patient (patient ‘PA’ as described above) whereas the remaining 5 groups all involved different patients. The three isolates from group G3 were collected within 6 weeks of each other (data not shown) even though the collection dates provided in this study indicate a 3 month period (Supplementary Table 1). The isolates for groups G1, G2, G5 and G6 were all collected within one month of each other.

### Antimicrobial resistance determinants of the G122 clade

Consistent with antibiotic susceptibility data for G122 isolates (5), the 94 NSW isolates had wildtype copies of the assessed AMR genes with the exception of harbouring the PBP2 345A insertion (Supplementary Table 4). Although the 345A insertion has been associated with penicillin-resistance (30) all NSW isolates were phenotypically susceptible to penicillin.

## Discussion

Overall the sequencing results show that isolates from the previously identified SNP-type are indeed closely related strains and, given they were all from females, provides further evidence of sustained transmission of this strain amongst heterosexuals. Moreover, the enhanced resolution provided by the phylogenetic analyses has provided new information regarding transmission of this strain within local sexual networks.

This study included six patients (PA to PF) for which more than one isolate was collected. The observed SNP variation within each patient was found to be minimal, ranging from 0-7 core SNPs. This within-patient genomic stability is not surprising and is consistent with a previous study (8). However, it provides an interesting context when considering the results for the five groups of patients (G1, G2, G3, G5 and G6; Figure 2) all of which involved different individuals having identical core SNPs. Such results are suggestive of tight transmission networks, perhaps even a common infection source for these patients. Of note were three females belonging to group G3 that shared identical core SNPs over a six week period. This observation would be consistent with a common infection source, possibly a male having multiple sexual partners and not actively seeking treatment. Behavioural data were not available to confirm this. Nevertheless, it should be noted that in the absence of these genomic data it would otherwise be very difficult to identify such clusters, particularly when patients are often unwilling or unable to provide details regarding sexual partners (31). Hence, these data highlight a potential role that WGS could have (assuming it could be achieved within a timely manner) in enhancing contact tracing, whereby the data are used to help pinpoint key groups of individuals for intervention.

The data also provide some evidence that CSW and certainly unprotected fellatio are not key drivers perpetuating this G122 genotype in the population. Previous reports have shown high rates of condom use for vaginal sex, but lower rates for both oral sex and non-paying partners among CSW (32, 33). Thus, if CSW were responsible then one would have expected clusters of pharyngeal infections among these individuals. In contrast, the phylogeny depicts clusters comprising a more even distribution of both pharyngeal and vaginal infections, suggesting that CSW alone is unlikely to be responsible for sustaining transmission of this genotype.

Of further interest in this study was the number of females in older age groups, with 21% of patients being 45 years or older. While there was some limited evidence of transmission within older ages (for example, cluster C2 predominately comprised individuals 45 years or older), the overall phylogeny suggested that gonorrhoea was not being sustained within distinct networks of older (≥ 45 years) individuals. Rather, the observed networks comprised a broad range of age groups and suggested that older individuals were acquiring infections that were otherwise being sustained in younger, more sexually active age groups. Further studies are needed to investigate this. However, current Australian federal policies promote sexual health to younger individuals and those of reproductive age as they are considered to be the most at risk for further complications (34). Given our data indicate networks of individuals comprising of varying ages, education on STI prevention may be more suited towards a broad range of age groups, including older ages and especially those engaging in new sexual relationships (35).

A recent comprehensive investigation applying WGS to *N. gonorrhoeae* isolates within Brighton, UK, allowed us to provide additional context for our genomes among a broader population. Interestingly, our common Australian strain only comprised 0.3% of isolates assessed from the Brighton study. It is difficult to speculate on the significance of these differences, however it may have only recently been introduced into the UK, perhaps via individuals travelling between the two regions. However, we note that NG-MAST 4186, which is associated with genotype G122, has been previously documented elsewhere, including Japan (36) and New Zealand (37) at relatively high prevalence (5.2% and 10.8% of strains respectively). Little is known as to why some *N. gonorrhoeae* strains may be more successful in a given population than others. Of interest was that our G122 genotype lacked almost all commonly reported *N. gonorrhoeae* AMR mechanisms, however current thinking is that these AMR mutations do not confer a fitness disadvantage (38). Therefore it is likely that other factors, possibly unrecognised virulence mechanisms, may be important in sustaining this genotype.

There were several limitations associated with this study, firstly that we did not access behavioural data; and secondly males were not assessed. However, the decision to exclude males was deliberate, and was done so as to focus on heterosexuals and limit any potential signal from MSM; Thirdly we only included isolates from two six-month time periods that were two years apart.

In summary, we have used whole genome sequencing to confirm transmission of a particular strain of *N. gonorrhoeae* amongst females in urban areas of Australia from 2012 and 2014. The use of WGS and its enhanced resolution has revealed features of local sexual networks that would not otherwise be apparent through routine surveillance activities. These data provide evidence of the additional value that WGS could provide in *N. gonorrhoeae* outbreak investigations. The information may also provide additional benefit for further studies aimed at identifying virulence markers that are important in maintaining such strains in the population.

## Acknowledgments

We thank Dr Tiffany Hogan from Prince of Wales Hospital, New South Wales for her assistance with this study. This study was conducted as part of the reference work of the National Neisseria Network (NNN), Australia, which is funded by the Australian Government Department of Health.

